# Recommendations for Population and Individual Diagnostic SNP Selection in Non-Model Species

**DOI:** 10.1101/2024.07.03.601943

**Authors:** Ellie E. Armstrong, Chenyang Li, Michael G. Campana, Tessa Ferrari, Joanna L. Kelley, Dmitri A. Petrov, Katherine A. Solari, Jazlyn A. Mooney

## Abstract

Despite substantial reductions in the cost of sequencing over the last decade, genetic panels remain relevant due to their cost-effectiveness and flexibility across a variety of sample types. In particular, single nucleotide polymorphism (SNP) panels are increasingly favored for conservation applications. SNP panels are often used because of their adaptability, effectiveness with low-quality samples, and cost-efficiency for use in population monitoring and forensics. However, the selection of diagnostic SNPs for population assignment and individual identification can be challenging. The consequences of poor SNP selection are under-powered panels, inaccurate results, and monetary loss. Here, we develop a novel user-friendly SNP selection pipeline for population assignment and individual identification, mPCRselect. mPCRselect allows any researcher, who has sufficient SNP-level data, to design a successful and cost-effective SNP panel for species of conservation concern.

## Introduction

Whole-genome sequencing (WGS) approaches have increased in feasibility and popularity as sequencing costs have declined and computational tools have improved. For many species, however, sequencing whole-genomes from a large number of individuals or using WGS for continuous population monitoring is still cost-prohibitive and computationally challenging. Well-established systems that rely heavily on genomic technologies, such as human or agricultural genetics, frequently turn to single-nucleotide polymorphism (SNP) panels or other forms of reduced-representation sequencing, such as RADseq or ddRAD for population assignment and sample identity.

SNP panels have been created for numerous non-model organisms including: Iberian lynx (Kleinman-Ruiz et al., 2017), puma (Fitak et al., 2015), lampreys (Hess et al., 2015), cichlids (Ciezarek et al., 2022), lions (Bertola et al., 2022), tigers (Natesh et al., 2019), bison (Wehrenberg et al., 2024), canids (Parker et al., 2022), and many others. This increase in popularity has occurred primarily because panels are cost-effective and flexible (Puckett, 2017). SNP panels are often designed to run on specific technologies (e.g. Fluidigm 96.96 Dynamic Array, Agena Bioscience MassARRAY), which limits them to specific facilities or ultimately requires additional validation on other machines (Carroll et al., 2018). Multiplex PCR (mPCR) is an alternative and more flexible approach, which involves creating a primer pool that amplifies many SNPs simultaneously. mPCR presents its own obstacles. For example, it can be challenging to avoid issues in the primer pools, such as primer dimers or cross amplification. However, because it is relatively straightforward to add indexes and adapters for many individuals and adapt primers to be compatible with a variety of sequencing technologies, mPCR also has considerable benefits. Methods such as GT-seq (Campbell et al., 2015) have made advances in the design and pooling of primers around specific loci, but the selection of those loci is left up to the user.

The selection of loci for a targeted panel is critical to ensure that panel aims (such as population assignment or sample identity) are accurately achieved. For example, it is possible to include SNPs that are not informative, which have the potential to introduce noise into downstream analyses. Further, when creating panels for multiple populations, SNPs that represent the genetic variation in one population may not be informative in the other due to allele frequency differences (Biddanda et al., 2020). This result has led the human genetics community to create multiple panels over time to better represent non-European human diversity, improve imputation accuracy, and bolster detection of variants that are associated with complex traits (Bien et al., 2016).

Panel design often includes the goals of population assignment and individual identification. The first hurdle in marker selection is that the methods used to filter and select SNPs that can achieve these two aims vary widely. Methods used for marker selection and filtering include but are not limited to, estimates of pi (**π**) or theta (θ), Hardy-Weinberg equilibrium (HWE), differentiation (*F*_ST_), minor allele frequency (MAF), linkage disequilibrium (LD), in conjunction with various quality filters such as mapping and genotype quality (see e.g. Fitak et al., 2015; Hess et al., 2015; Kleinman-Ruiz et al., 2017; Natesh et al., 2019; Bertola et al., 2022; Ciezarek et al., 2022; Wehrenberg et al., 2024). Human geneticists previously developed approaches to select optimal genetic markers for ancestry (Balding & Nichols, 1994; Rosenberg et al., 2003; Rosenberg, 2005; Manel et al., 2005; Kidd et al., 2006; Baye et al., 2009; Galanter et al., 2012) and individual identification (Balding & Nichols, 1994; Kidd et al., 2006; Pakstis et al., 2007, 2010). However, these insights are not always applied in non-model species. Significantly, a majority of the theory that guides SNP selection for population assignment and individual identification assumes that markers are in linkage equilibrium and segregating at an appreciable frequency within populations, making MAF and LD filtering most relevant during the SNP filtering stage (Kidd et al., 2006). Further, in human genetics, filtering for HWE is often used to ensure genotype quality since HWE outlier loci are often caused by sequencing errors. Whereas in conservation genetics, HWE filters have been shown to remove informative loci that are variable between populations (Chen et al., 2017; Pearman et al., 2022), primarily because conservation projects have substantially lower sample sizes. Given the numerous approaches that exist and the potential large variance of their success in non-human species, the selection of loci remains challenging for conservation-oriented projects.

Adding to this list of considerations for marker selection, researchers must also assess marker informativeness to build robust SNP panels. Once again, there are many different statistics that measure marker informativeness from the perspective of population assignment, including Fisher Information Content (FIC; Pfaff et al., 2004), Shannon Information Content (SIC; Rosenberg et al., 2003), F statistics (in particular, *F*_ST_ ; Wright, 1951), Informativeness for Assignment Measure (I_n_; Rosenberg et al., 2003), and the Absolute Allele Frequency Differences (delta, δ; Rosenberg et al., 2003). Principal Component Analysis (PCA) has additionally been used as a tool to select SNPs for structure identification and assignment (Paschou et al., 2007). Previous work has suggested that the two best methods of estimating marker informativeness for biallelic loci are I_n_ and *F*_ST_ (Ding et al., 2011), with I_n_ performing better for mixed ancestry populations. Studies have also shown that selecting markers which maximize *F*_ST_ perform better than selecting markers using PCA: (Wilkinson et al., 2011).

In addition to estimating individual marker informativeness, there are methods for determining which combinations of markers will yield the most effective panel, such as *f*_ORCA_ (Rosenberg, 2005). This approach has rarely been implemented outside of human and agriculturally-relevant species, such as salmon (Storer et al., 2012), sheep (Sottile et al., 2018), and crop species (Morrell & Clegg, 2007). In rare cases, the method has been applied to non-model systems, such as in the domestic cat and European wildcat (Oliveira et al., 2015). Despite its relevance to optimizing the selection of small panels (which are desirable in the conservation sector), this method has seldom been applied in non-model species.

Here, we seek to optimize diagnostic SNP selection using *f*_ORCA_, specifically for the purposes of population assignment. We demonstrate the applicability of these limits in humans, tigers, and domestic dogs. We also briefly explore the overlap between selecting SNPs for population assignment and individual identification. Last, we present an accompanying pipeline, mPCRselect, that when provided with a variant call file (VCF) and pre-designated populations uses a greedy algorithm to provide users with diagnostic SNPs suitable for population assignment and individual identification in the context of a multiplex PCR assay.

## Materials and Methods

### Population assignment performance function

We assigned individuals to populations with a performance function, *f*_ORCA_ (Rosenberg et al., 2003), which uses the genotype of an individual and population-level allele frequencies of the selected marker for population assignment. For a detailed description of the *f*_ORCA_ approach see (Rosenberg, 2005). Briefly, for each sample, we calculated the probability that the individual originated from a given source population (k), then we assigned the individual to the population that had the highest probability.

### Simulated genotype data

We conducted forward-in-time simulations using SLiM 4.0.1 (Haller & Messer, 2023) on a high-performance computing cluster, utilizing Dell node models R440 and xl170 with core speeds of 2.1Ghz. We simulated a burn-in with neutral dynamics for a population of 10,000 individuals, each with a uniform 100Mb genomic segment, until all lineages in the ancestral trees were fully coalesced. After the burn-in, the ancestral population was instantaneously split into two subpopulations of 10,000 individuals each. We allowed no migration between subpopulations after divergence. Simulations were conducted with a neutral mutation rate of 1e-8 and a recombination rate of 1e-8. As the subpopulations diverged, we recorded individual genotypes every 200 generations until 2000 generations after the population split. Increments of 200 generations were selected based on theoretical estimates of *F*_ST_ from (Nicholson et al., 2002), and resulted in the populations having an *F*_ST_ ranging [0.01, 0.1] with increments of 0.01.

Finally, we sub-sampled 100 individuals from each of the two populations and output a VCF for classification. The VCF was converted to a plink file with PLINK 2 (version 2.00a3.7LM; (Chang et al., 2015)). The plink file was filtered to 10,000 markers in linkage equilibrium with each other with the command ‘-indep-pairwise 500kb 0.2’.

### Empirical Data

The marker selection method was applied to three different datasets. We used whole-genome sequence data from humans in the 1000 Genomes Project (Dai, CDX; Puerto Rican, PUR; Luhya, LWK; Colombian, CLM; and Afro-Caribbean, ACB) (1000 Genomes Project Consortium et al., 2010); from tigers (Amur, Bengal, and Generic, Armstrong et al., 2024); and dogs (Labrador retriever and Yorkshire terriers, Mooney et al., 2023). Sample sizes were 88 humans from each population (N = 440 individuals total), 13 tigers from each of the Amur and Bengal subspecies, and 13 Generic (N = 39 individuals total), and 100 individuals from each dog breed (N = 200 individuals total). Unrelated individuals from the 1000 Genomes Project were identified using the ped file (20130606_g1k.ped) that is provided with the hg19 data. For the tiger data, unrelated individuals were previously identified in Armstrong et al., 2024. For the dog data, unrelated individuals were previously identified in Mooney et al., 2021. We used PLINK 2 (version 2.00a3.7LM; Chang et al., 2015) and filtered for a minor allele frequency that was at least 5% (common in the population) and markers that were in linkage equilibrium with each other. We used the command ‘--maf 0.05 -indep-pairwise 500kb 0.2’ to accomplish this filtering. Then, we created a subset of 10,000 markers randomly sampled from across the genome using the remaining markers in linkage equilibrium.

### Marker Selection and Population Assignment

*F*_ST_ (Wright, 1951) was used to quantify the genetic differentiation among subpopulations. The higher the *F*_ST_ value, the more pronounced the genetic differentiation. Various definitions and formulations of *F*_ST_ exist, and in this work we use

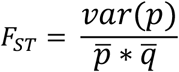

to calculate *F*_ST_ values (Balding, 2003). We calculated *F*_ST_ values for all of the markers and sorted the results in descending order. A greedy algorithm was applied to select the top M markers to compose a marker panel for individual classification.

Population-level allele frequencies were computed on the basis of the sampled individuals. We used *f*_ORCA_ to assign N individuals a population label computed with either M random markers or the M top markers. Accuracy was measured as the proportion of empirical individuals that were classified with the correct population label. We ran 20 replicates of each classification scenario in the simulated and empirical datasets.

### mPCRselect pipeline

We designed a novel Nextflow pipeline to select optimal SNP markers for population differentiation and individual identification. The pipeline is compatible with macOS and Linux operating systems and can run locally on a desktop/laptop computer or remotely on a computing cluster or in the cloud. Most dependencies can be installed automatically in their own Conda environment through Nextflow (Di Tommaso et al., 2017). Only the optional dependencies (BaitsTools; Campana, 2018 and NGS-PrimerPlex; Kechin et al., 2020) must be installed by the end-user as no Conda recipe currently exists for these packages. We describe the pipeline briefly below. A simplified flow diagram of the pipeline is available as Supplementary Figure. See the mPCRselect documentation (https://github.com/ellieearmstrong/mPCRselect) for a complete diagram including the various parameters, internal processes, and outputs.

mPCRselect’s primary input files are a variant call format (VCF) file of individual genotypes and a comma-separated value (CSV) table assigning each individual in the VCF to a designated population. Optional input includes a list of individuals to remove from the dataset, a list of chromosomes to retain in the analysis, and a browser extensible data (BED) format file of genomic coordinates to remove from the dataset (e.g. regions of low mappability, repetitive regions, etc.). mPCRselect uses VCFtools (Danecek et al., 2011) to remove unwanted individuals, chromosomes, and genomic regions from the dataset. VCFtools is also used to identify and remove singleton and individual-unique doubleton sites, remove sites that failed previously applied filters (e.g. those without a ‘PASS’ flag’), exclude non-biallelic sites, and to filter the input VCF by genotype quality (GQ) and site and individual data missingness. The pipeline uses a custom Python script (‘Culling.py’) to remove SNPs that are within a user-specified distance of another SNP (default is 40 bp; recommended based on typical size of amplicons). This filter aims to exclude variants where primer attachment sites would contain additional SNPs and inhibit amplification. Afterwards, the sites are optionally thinned by physical distance using VCFtools and by LD using PLINK 2 (Chang et al., 2015). The resulting file then is pushed through two distinct paths to select markers for population assignment and markers for individual identification, respectively.

For population assignment, we implement the greedy algorithm described above using a custom R script (‘make_fst_plots.R’). In order to account for differences in sample size between populations, we bootstrap individuals to a user-specified population size (default is 20 individuals per population). The greedy algorithm is run a user-specified number of times (default is 20 repetitions per comparison of two populations). Users are provided with output plots which correlate the number of markers with assignment accuracy for each repetition. Additionally, mPCRselect identifies the sites with the highest *F*_ST_ values overall using VCFtools (flag ‘--weir-fst-pop’). mPCRselect then uses a custom Ruby script (‘get_best_snps.rb’) to identify the sites that appear most frequently in the lists of highest *F*_ST_ sites from each of the greedy algorithm runs and from the VCFtools *F*_ST_ analysis.

To select markers for individual identification, the VCF file is first split into distinct populations and **π** is then estimated using VCFtools with the flag ‘--site-pi’, which calculates nucleotide divergence on a per site basis using the following equation:

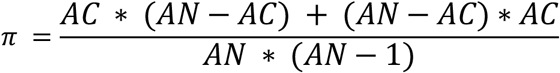

where AC = allele count and AN = allele number. We use **π** in order to select sites for individual identification because **π** will also be maximized at alleles with intermediate frequencies. For each population, we compile a list of sites with the highest **π**. We then use the ‘get_best_snps.rb’ script to identify the most frequently appearing sites across the datasets. Afterwards, we calculate the random match probability (RMP) of the selected SNPs using a custom R script (‘RMP_calc.R’).

After site selection, the individual identification and ancestry assignment SNP sets are combined and the user can choose to design baits for capture-based projects using BaitsTools (Campana, 2018) or alternatively import the sites into NGS-PrimerPlex (Kechin et al., 2020) to design primers for multiplex SNP panels. Finally, as an *in silico* validation of the panel design, the program performs PCA (using PLINK 2) on the biallelic sites from the unfiltered input VCF, the post-filtering VCF, and the final chosen population-assignment and individual-identification sites (both separately and concatenated).

## Results

### Population assignment using realistic simulated data

To better understand how *F*_ST_ influenced the ability of *f*_ORCA_ to assign individuals to a population, we simulated a split-model of two populations with no migration in SLiM. During classification, we also varied the number of individuals (N) and number of markers (M) being used for population assignment. Then, for each set of parameters, we computed the rate of misclassification while varying N, M, and *F*_ST_ values. Overall, we found that when *F*_ST_ between populations is 0.01, individuals could be correctly assigned to a population with as few as 10 markers (Figure 1), though the probability of incorrectly assigning individuals is quite large, the average misclassification rate was approximately 37% (Supplementary Table 1). Importantly, increasing the marker set to as few as 100 markers decreased the average misclassification rate to approximately 12% (Supplementary Table 2). Conversely, when *F*_ST_ = 0.09, which is on the same order of magnitude as the average *F*_ST_ between two human populations (Ramachandran et al., 2005; Rosenberg et al., 2005), 10 markers resulted in an average misclassification rate of 21% (Supplementary Table 1). When we increased the marker set and included 80 markers, we achieved misclassification rates that were on-average less than 1% (Figure 1). Increasing the number of markers improved accuracy for every tested value of *F*_ST_ (Figure 1). Our results demonstrate that when *F*_ST_ is small (0.01), it is necessary to include more markers for accurate classification (Supplementary Tables 1 & 2), and as *F*_ST_ increases accurate results can be achieved with fewer markers and fewer individuals (Figure 1).

**Figure 1:**
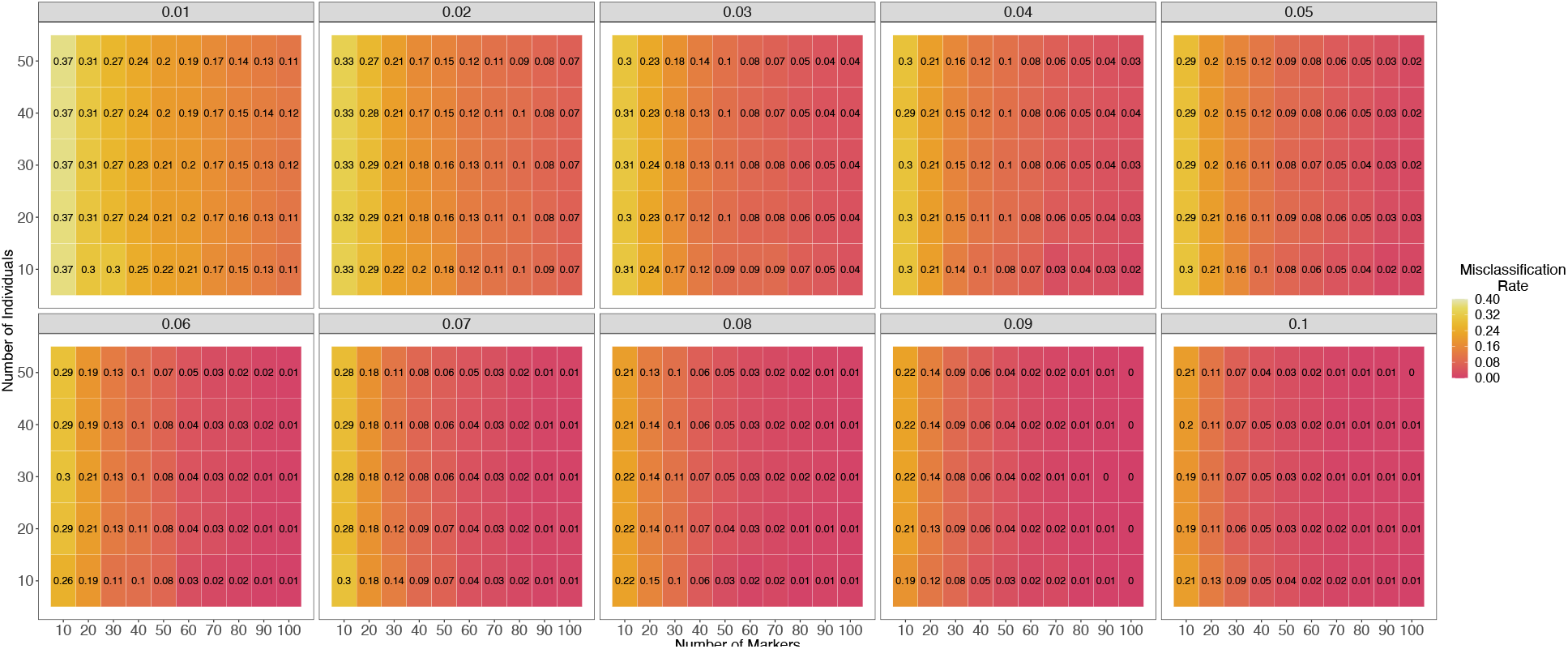
Classification using simulated data. Two populations with a given measure of *F*_ST_ (as indicated in the gray bar above each panel) were classified using (M) markers (x-axis) and that consisted of (N) individuals per-population (y-axis). The misclassification rate ranges from 0 (pink) to 0.4 (light yellow). As the *F*_ST_ value and number of markers increases, the misclassification rate decreases.

### Empirical Data

After testing the algorithm with simulated data, we tested the method using three empirical data sets from humans, tigers, and domestic dogs. First, we compared publicly available human data from the 1000 Genomes Project (1000 Genomes Project Consortium et al., 2010). We used whole-genome sequence data from an African population with a single origin, the Luhya from Kenya; and an Asian population with a single origin, the Dai from China (Figure 2A). When the number of markers was the smallest, we observed the largest accuracy gain using the markers selected from the method developed here versus randomly selected markers. For example, when the number of markers was the smallest (M=10), the average accuracy across random marker sets was 0.8884±0.0651, while the average accuracy across top marker sets is a perfect accuracy of 1. When the number of randomly selected markers was twenty, we observed an average accuracy across groups of 0.9611±0.0040, compared to the top marker set which had an average accuracy of 1. As the number of markers increased, the accuracy gain decreased. This pattern was observed across all data sets.

**Figure 2:**
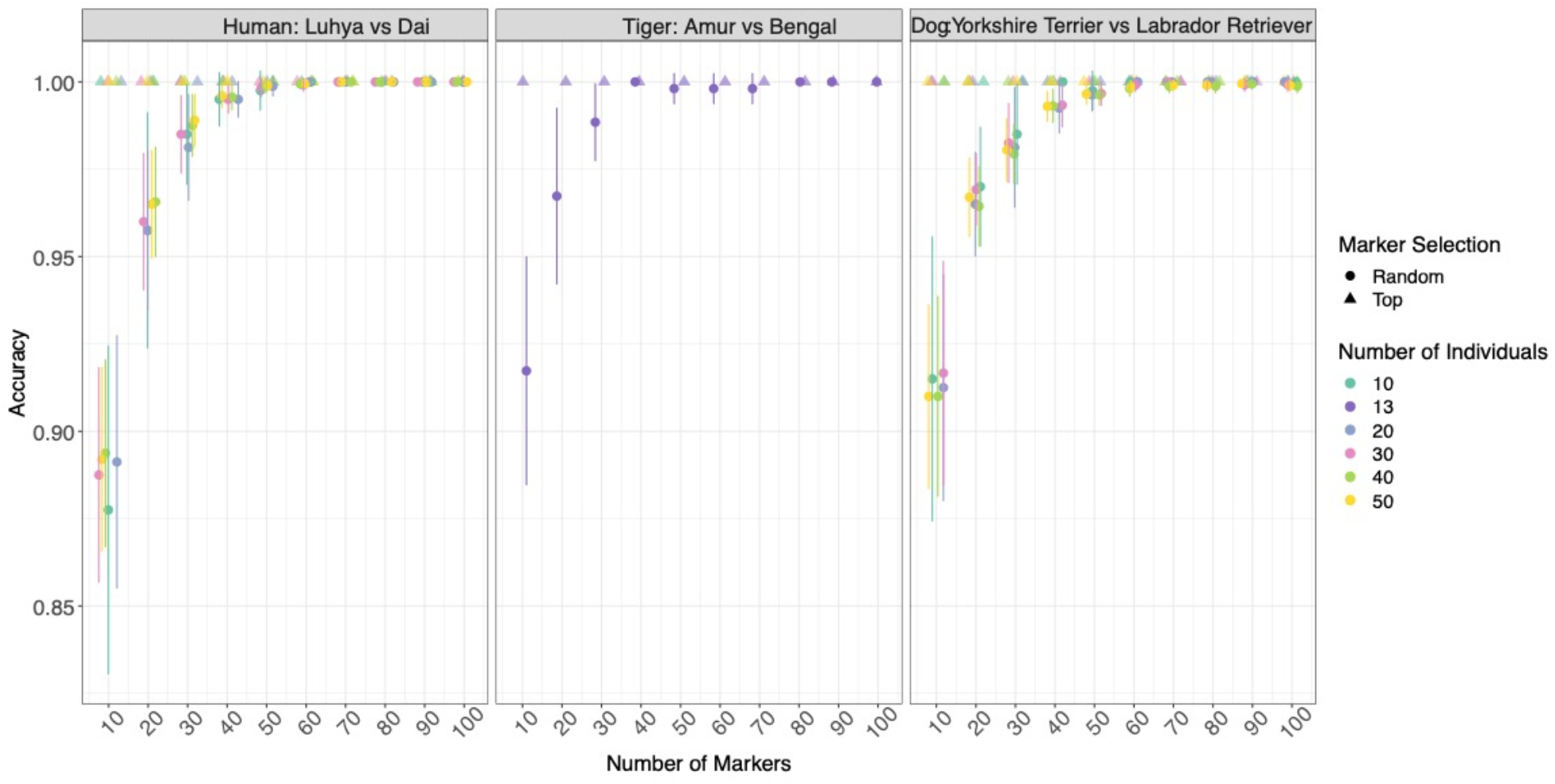
Classification accuracy using two approaches, 1) top *F*_ST_ markers (triangles) and 2) random markers (circles). For the random markers, each dot signifies the mean over 20 simulation replicates. The x-axis indicates the number of markers used for classification and the dot color indicates the number of individuals. The accuracy of classification is shown on the y-axis. Here, three classifications were conducted using human populations (Luhya and Dai), tiger populations (Amur and Bengal), and breed dogs (Yorkshire Terrier and Labrador Retriever).

Given their endangered status and relatively recent efforts to sequence tiger populations, we only had access to 13 Amur individuals for performing classification (Figure 2B). The limited sample size also impacted our ability to vary the number of individuals we sampled for classification. However, because the *F*_ST_ value (∼0.2; Armstrong et al., 2021) between the wild Amur and Bengal tiger populations is larger than that of human populations, even using only 10 random markers, the accuracy of classification is 0.9173±0.0652 (Figure 2A & B). This accuracy was higher than what was achieved in the human dataset with the same number of markers for each set of classifications. Additionally, the random marker set began to match the top markers more quickly in the tigers than humans.

Lastly, we examined breed dogs (Figure 2C), which have high homozygosity within breeds, but are quite divergent between the various clades (Parker et al., 2017). Since both dog and tiger populations had larger *F*_ST_ than the human populations, we once again observed that the random marker set started with a high classification accuracy (0.9128±0.0639), and reached the same performance as top markers faster relative to humans (Figure 2).

We also created a dataset that allowed us to explore whether the degree of admixture influenced our ability to accurately classify populations (Figures S1 & S2). In order to achieve this, we conducted classification in additional human populations, specifically Puerto Rican, Colombian and Afro-Caribbean, (Figure S1) and the captive (Generic) tiger population (Figure S2). We found that admixture decreased classification accuracy in both human and tiger populations. The drop in accuracy was most impactful when using random marker sets (Figures S1 & S2) and less severe when using *f*_ORCA_ in conjunction with the algorithm developed here which selected the top *F*_ST_ markers. It is also important to note that the degree to which there was shared ancestry between the two populations played a role in the magnitude of the decreased accuracy. For example, the Puerto Rican population represented in 1000 genomes has a more similar ancestry composition to sampled Colombian population than the sampled Afro-Caribbean population (1000 Genomes Project Consortium et al., 2010). Thus, the classification is much worse across all random marker sets in the Colombian population compared with the Afro-Caribbean. Classification also required more top *F*_ST_ markers to perform accurately in the admixed populations. In the tigers, we observed a similar pattern when comparing classification accuracy for differentiating the Amur and Bengal populations versus the Amur and Generic (captive) populations. On average, a random individual in the captive population could have up to approximately 39% Amur ancestry (Armstrong et al., 2024), which led to a marked (0.9173±0.0652 to 0.8442±0.0813) drop in our classification accuracy at the minimum marker set, M=10, and a lag in the random marker classification accuracy reaching the accuracy of the top markers.

Overall, our method always outperformed the randomly selected marker sets. In order to accurately classify population pairs with lower *F*_ST_ values, we always had to use more markers. For randomly selected markers, as more markers are incorporated into the panel there was consistently better performance for population assignment. This is similar to previous results from both theory and empirical data (McVean, 2009; Patterson et al., 2006; Rosenberg, 2005).

### Allele frequency differences of the most informative markers

We next examined the allele frequency distributions of the top marker sets in humans, tigers, and dogs (Figure 3). As we expected, the majority of top *F*_ST_ markers were concentrated at opposite allele frequencies, irrespective of the compared populations. Given that *F*_ST_ quantifies the magnitude of drift between two populations, this was expected. The allele frequency differences were smallest in the human populations studied (Figure 3A) and largest in tigers and dogs (Figure 3B & C), which have a larger value of *F*_ST_ overall.

**Figure 3:**
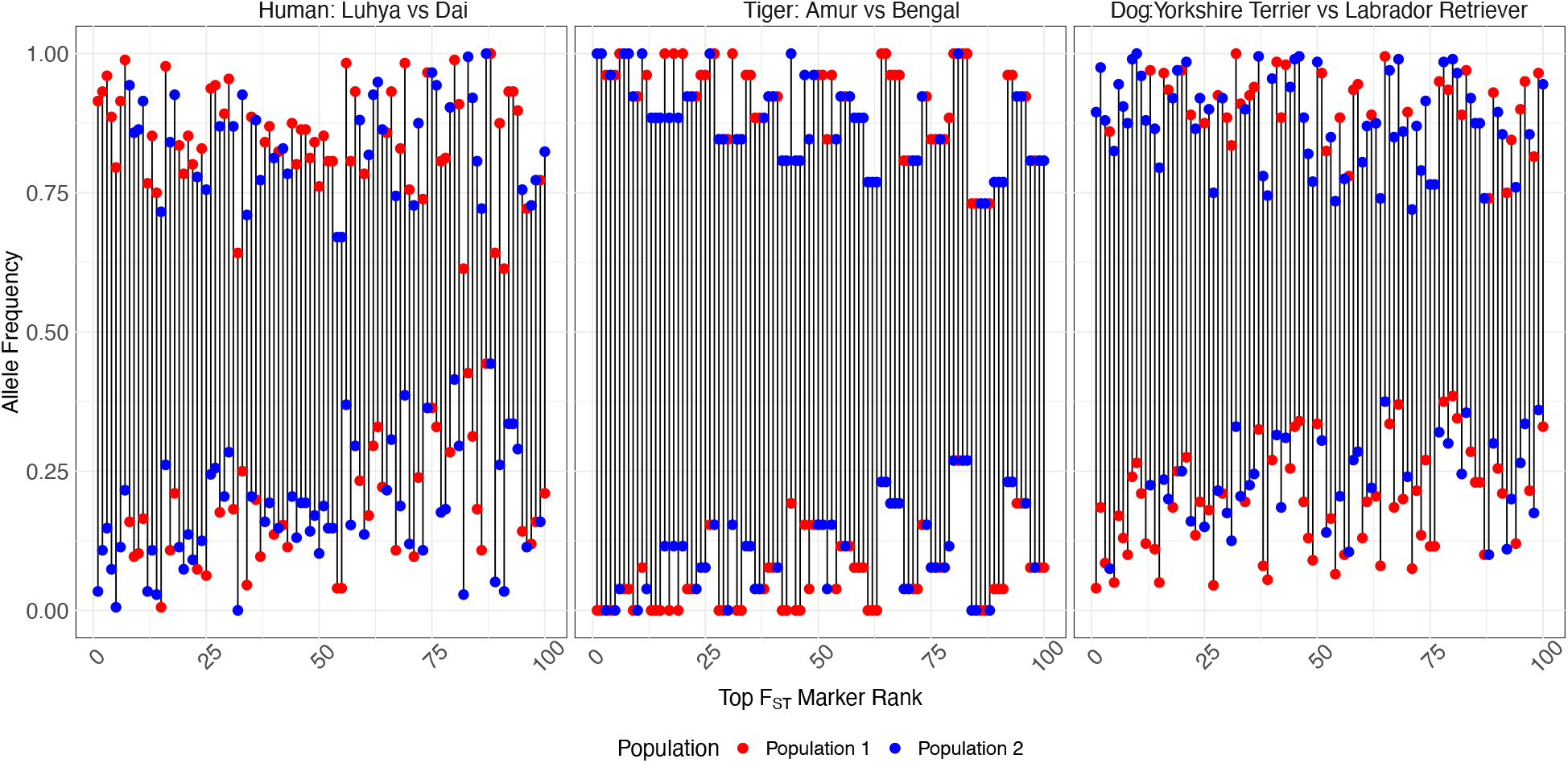
Allele frequency of the top 100 *F*_ST_ markers between populations and the frequency difference for each marker between various populations. The x-axis indicates the rank of *F*_ST_ values of the markers, and the y-axis indicates the markers’ allele frequency in both Luhya (represented in blue) and Dai (represented in red) populations; and Amur (represented in blue) and Bengal (represented in red); Labrador Retriever (represented in blue) and Yorkshire Terrier (represented in red).

Taking the same approach as with classification, we explored whether the degree of admixture influenced the magnitude of the allele frequency gap of top *F*_ST_ markers between the same admixed human and captive (Generic) tiger populations (Figures S3 & S4). Indeed, admixture decreased the allele frequency gap and that the degree to which there was shared ancestry affected how close the allele-frequency gap was. This was expected, given that recent shared ancestry will likely result in populations having more similar allele frequencies (Ramachandran et al., 2005). Since the Puerto Rican population has a more similar ancestry composition to sampled Colombian population, we expected that the allele frequency gap would be smaller than both the Afro-Caribbean and Luhya populations, which was ultimately what we observed (Figure S3). The tigers followed this expectation as well, and the allele frequency gap between the Amur and Bengal tigers was larger than that between the Amur and Generic tigers (Figure S4).

### The relationship between top ancestry markers and random match probability

One measure of the degree to which a marker is useful in individual identification is the probability that two random individuals from the population have matching genotypes at the marker—if this probability is low, then the marker will distinguish individuals often. We will refer to this probability as the Random Match Probability (RMP). Note, we are using the term differently than it is defined in forensics, in forensics RMP refers to the probability that a random person matches a pre-specified genotype, rather than the probability that two randomly sampled individuals match at a given locus. For biallelic loci at Hardy-Weinberg equilibrium, this probability is:

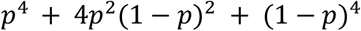

The equation above is minimized at allele frequency *p* = 0.5. In other words, markers with allele frequencies near 0.5 are the most useful for individual identification. Such markers tend to not be found in our top *F*_ST_ marker sets (Figure 3). In tigers and dogs, markers with high *F*_ST_ seldom had allele frequencies near 0.5 in either population. For human populations, though we found more alleles that existed at frequencies closer to 0.5 we still did not observe a strong overlap between top *F*_ST_ and RMP markers when comparing our reference population. When we identified the top 100 markers for minimizing RMP, we observed no overlapping markers in the Luhya population, no overlapping markers in the Amur subspecies, and no overlapping markers in Yorkshire Terriers when the marker set was overlapped with the top *F*_ST_ markers in Figure 3. When we further explored the overlap between top *F*_ST_ and RMP markers, we found that two markers overlapped when comparing the Puerto Rican and Luhya populations, one marker overlapped when comparing the Puerto Rican and Colombian populations, two markers overlapped when comparing the Puerto Rican and Afro-Caribbean populations, and no markers overlapped when comparing the Generic and Amur populations.

## Discussion

This work introduces the mPCRselect pipeline which is designed to provide users with a sufficient marker set to distinguish populations within their species of interest and/or a marker set to identify individuals. Markers for population assignment are selected using *F*_ST_ and tend to be close to fixation or loss when comparing two populations, while markers which optimize individual identification hold intermediate frequencies within populations. Implementing our marker selection method consistently reduces the number of markers required for accurate population assignment.

Our findings are consistent with previous research, where the relationship between the number of SNPs, their frequency, and the power to detect differentiation between populations has previously been explored in a conservation context (Morin et al., 2009; Willing et al., 2012). Morin et al. 2009 primarily explored this relationship from the perspective of initial study design (i.e. determining how many loci are necessary to detect population structure without having knowledge of the MAF or linkage status of a marker) and confirmed that more SNPs are necessary to detect differentiation between groups with lower values of *F*_ST_. Willing et al. 2012 also echoed results from (Patterson et al., 2006) showing that even with small sample sizes, large numbers of markers can compensate to provide accurate estimates of *F*_ST_. A graphical user interface with the explicit goal of helping conservation practitioners select the appropriate number of samples and markers and avoid suboptimal sampling was presented in (Hoban et al., 2013), again from the perspective of initial study design. Critically, for each study, a sufficient number of individuals must be sampled in order to get an accurate estimate of allele frequencies in the population and *F*_ST_. Our method is in-line with these findings, but we approach the other end of the problem when populations have already been identified and one desires identifying optimal markers. We found that population assignment accuracy increases as more informative markers are added. We observed the lowest accuracy when we used the smallest set of random markers (M=10), and as the number of random markers increased they achieved a performance similar to the top markers. This limitation on information content when using *F*_ST_ for marker selection was previously highlighted in several studies (e.g Balding & Nichols, 1994; Rosenberg et al., 2003; Rosenberg, 2005; Manel et al., 2005; Kidd et al., 2006; Baye et al., 2009; Galanter et al., 2012). Our method, which identifies the most informative markers by computing *F*_ST_, then conducts population assignment with f_ORCA_ consistently outperforms or does as well as random markers when a sufficient set size is achieved. However, we emphasize the findings of previous studies which show that sufficient data is necessary for detecting population structure initially (e.g. that sufficient markers and individuals are required to detect structure between populations; Patterson et al. 2006, Morin et al. 2009, Willing et al. 2012). Insufficient data at this stage would ultimately result in inaccurate allele frequencies and hinder the accurate estimation of *F*_ST_ and **π**, which are critical for optimal marker selection.

It is important to note that populations with higher divergence (as measured by *F*_ST_) will require fewer markers for classification, whereas populations that are less divergent will require more markers. In the future, one natural extension to explore is optimizing markers for RMP and population assignment simultaneously, rather than as distinct steps. Incorporating RMP will tend to increase the number of markers on the panel, because markers that are most informative for population classification (Figure 3) typically have relatively low minor allele frequencies in each subpopulation. Conversely, RMP is maximized by alleles of intermediate frequency. Thus, for biallelic loci, the top *F*_ST_ markers, which are the markers most useful in population assignment, tend to be the least useful markers in the individual identification. This contrasts with the situation for multi-allelic human STRs, where a correlation has been observed between markers’ usefulness for individual identification and for population classification (Algee-Hewitt et al., 2016). Another natural extension of this method could be for identification and selection of marker sets for sex identification and relatedness.

Though we do not discuss the practical applications of applying SNP panels here, it is broadly recommended to design and test more primer sets than what is identified as the ‘minimum number’ to achieve successful population assignment or individual identification, since some SNPs will not amplify as well as others, or may not work due to other factors such as primer dimerization. Panels designed for low-quality samples (scat, hair, environmental, or forensic materials) may require an increase in the overall number of SNPs being screened, as drop out will occur due to sample degradation, which will impact the ability to accurately assign or identify any one sample. This must be balanced with the expected amount of endogenous DNA in the sample because primers will not amplify well if there is too little DNA template.

Our novel pipeline mPCRselect streamlines SNP panel design for ancestry assignment and individual identification. Further, the flexibility of this software allows for straight-forward integration of novel algorithms for marker selection in the future. Similar pipelines have already been created for selecting markers from human genomic data (S. Chen et al., 2020). One such pipeline is the R Package AIMsetfinder, which also uses a Bayesian approach to identify SNPs which are informative of population assignment. However, their approach is different from the approach described here because it tests every marker in the dataset, then goes backwards to create a set of markers that minimize a logarithmic loss function (Pfaffelhuber et al., 2020). Contrastingly, our pipeline maximizes our assignment function, f_ORCA_, using the most informative markers while simultaneously providing the user with a seamless connection to primer or amplicon design software.

In sum, to create an effective SNP panel, one must carefully consider the minimum number of markers that should be present on the panel and the number of individuals to sample. These values should be determined by how divergent the populations of interest are and how accurate the classification needs to be. Our pipeline mPCRselect streamlines the process of selecting optimized markers for population assignment and individual identification for any user with sufficient data.

## Acknowledgements

We thank Michael “Doc” Edge for extremely thoughtful and helpful comments and conversations about this work. E. E. Armstrong was supported by a Washington Research Foundation Postdoctoral Fellowship. C.L. was supported by the WiSE Summer Undergraduate Research Fellowship. J.A.M was supported by the startup funds from Dornsife College of Letters, Arts and Sciences through the Department of Quantitative and Computational Biology and the USC WiSE Gabilan Assistant Professorship. M.G. Campana was supported by the Smithsonian Institution.

## Data accessibility and benefit-sharing

Summary files, simulation scripts, classification code, and empirical data for this project is provided at GitHub repository https://github.com/ChenyangLi6/SNP-panel. mPCRSelect is available at https://github.com/ellieearmstrong/mPCRselect.

## Author contributions

E.E.A. and J.A.M. conceived of the project. C.L., J.A.M., and T.F. performed simulations and data analysis. E.E.A., C.L., and M.G.C. built the mPCRselect pipeline, with contributions from K.A.S. and J.A.M. E.E.A., C.L., and J.A.M. wrote the manuscript. M.G.C, J.L.K., D.A. P., and K.A.S., provided feedback and input on manuscript content and analyses. All authors approved of the final manuscript.

## References

1000 Genomes Project Consortium, Abecasis, G. R., Altshuler, D., Auton, A., Brooks, L. D., Durbin, R. M., Gibbs, R. A., Hurles, M. E., & McVean, G. A. (2010). A map of human genome variation from population-scale sequencing. Nature, 467(7319), 1061–1073.

Algee-Hewitt, B. F. B., Edge, M. D., Kim, J., Li, J. Z., & Rosenberg, N. A. (2016). Individual Identifiability Predicts Population Identifiability in Forensic Microsatellite Markers. Current Biology: CB, 26(7), 935–942.

Armstrong, E. E., Khan, A., Taylor, R. W., Gouy, A., Greenbaum, G., Thiéry, A., Kang, J. T., Redondo, S. A., Prost, S., Barsh, G., Kaelin, C., Phalke, S., Chugani, A., Gilbert, M., Miquelle, D., Zachariah, A., Borthakur, U., Reddy, A., Louis, E., … Ramakrishnan, U. (2021). Recent Evolutionary History of Tigers Highlights Contrasting Roles of Genetic Drift and Selection. Molecular Biology and Evolution, 38(6), 2366–2379.

Armstrong, E. E., Mooney, J. A., Solari, K. A., Kim, B. Y., Barsh, G. S., Grant, V. B., Greenbaum, G., Kaelin, C. B., Panchenko, K., Pickrell, J. K., Rosenberg, N., Ryder, O. A., Yokoyama, T., Ramakrishnan, U., Petrov, D. A., & Hadly, E. A. (2024). Unraveling the Genomic Diversity and Admixture History of Captive Tigers in the United States. In bioRxiv (p. 2023.06.19.545608). 10.1101/2023.06.19.545608

Balding, D. J. (2003). Likelihood-based inference for genetic correlation coefficients. Theoretical Population Biology, 63(3), 221–230.

Balding, D. J., & Nichols, R. A. (1994). DNA profile match probability calculation: how to allow for population stratification, relatedness, database selection and single bands. Forensic Science International, 64(2-3), 125–140.

Baye, T. M., Tiwari, H. K., Allison, D. B., & Go, R. C. (2009). Database mining for selection of SNP markers useful in admixture mapping. BioData Mining, 2(1), 1.

Bertola, L. D., Vermaat, M., Lesilau, F., Chege, M., Tumenta, P. N., Sogbohossou, E. A., Schaap, O. D., Bauer, H., Patterson, B. D., White, P. A., de Iongh, H. H., Laros, J. F. J., & Vrieling, K. (2022). Whole genome sequencing and the application of a SNP panel reveal primary evolutionary lineages and genomic variation in the lion (Panthera leo). BMC Genomics, 23(1), 321.

Biddanda, A., Rice, D. P., & Novembre, J. (2020). A variant-centric perspective on geographic patterns of human allele frequency variation. eLife, 9. 10.7554/eLife.60107

Bien, S. A., Wojcik, G. L., Zubair, N., Gignoux, C. R., Martin, A. R., Kocarnik, J. M., Martin, L. W., Buyske, S., Haessler, J., Walker, R. W., Cheng, I., Graff, M., Xia, L., Franceschini, N., Matise, T., James, R., Hindorff, L., Le Marchand, L., North, K. E., … PAGE Study. (2016). Strategies for Enriching Variant Coverage in Candidate Disease Loci on a Multiethnic Genotyping Array. PloS One, 11(12), e0167758.

Campana, M. G. (2018). BaitsTools: Software for hybridization capture bait design. Molecular Ecology Resources, 18(2), 356–361.

Campbell, N. R., Harmon, S. A., & Narum, S. R. (2015). Genotyping-in-Thousands by sequencing (GT-seq): A cost effective SNP genotyping method based on custom amplicon sequencing. Molecular Ecology Resources, 15(4), 855–867.

Carroll, E. L., Bruford, M. W., DeWoody, J. A., Leroy, G., Strand, A., Waits, L., & Wang, J. (2018). Genetic and genomic monitoring with minimally invasive sampling methods. Evolutionary Applications, 11(7), 1094–1119.

Chang, C. C., Chow, C. C., Tellier, L. C., Vattikuti, S., Purcell, S. M., & Lee, J. J. (2015). Second-generation PLINK: rising to the challenge of larger and richer datasets. GigaScience, 4, 7.

Chen, B., Cole, J. W., & Grond-Ginsbach, C. (2017). Departure from Hardy Weinberg Equilibrium and Genotyping Error. Frontiers in Genetics, 8, 167.

Chen, S., Ghandikota, S., Gautam, Y., & Mersha, T. B. (2020). MI-MAAP: marker informativeness for multi-ancestry admixed populations. BMC Bioinformatics, 21(1), 131.

Ciezarek, A., Ford, A. G. P., Etherington, G. J., Kasozi, N., Malinsky, M., Mehta, T. K., Penso-Dolfin, L., Ngatunga, B. P., Shechonge, A., Tamatamah, R., Haerty, W., Di Palma, F., Genner, M. J., & Turner, G. F. (2022). Whole genome resequencing data enables a targeted SNP panel for conservation and aquaculture of Oreochromis cichlid fishes. Aquaculture, 548, 737637.

Danecek, P., Auton, A., Abecasis, G., Albers, C. A., Banks, E., DePristo, M. A., Handsaker, R. E., Lunter, G., Marth, G. T., Sherry, S. T., McVean, G., Durbin, R., & 1000 Genomes Project Analysis Group. (2011). The variant call format and VCFtools. In Bioinformatics (Vol. 27, Issue 15, pp. 2156–2158). 10.1093/bioinformatics/btr330

Ding, L., Wiener, H., Abebe, T., Altaye, M., Go, R. C. P., Kercsmar, C., Grabowski, G., Martin, L. J., Khurana Hershey, G. K., Chakorborty, R., & Baye, T. M. (2011). Comparison of measures of marker informativeness for ancestry and admixture mapping. BMC Genomics, 12, 622.

Di Tommaso, P., Chatzou, M., Floden, E. W., Barja, P. P., Palumbo, E., & Notredame, C. (2017). Nextflow enables reproducible computational workflows. Nature Biotechnology, 35(4), 316–319.

Fitak, R. R., Naidu, A., Thompson, R. W., & Culver, M. (2015). A New Panel of SNP Markers for the Individual Identification of North American Pumas. Journal of Fish and Wildlife Management, 7(1), 13–27.

Galanter, J. M., Fernandez-Lopez, J. C., Gignoux, C. R., Barnholtz-Sloan, J., Fernandez-Rozadilla, C., Via, M., Hidalgo-Miranda, A., Contreras, A. V., Figueroa, L. U., Raska, P., Jimenez-Sanchez, G., Zolezzi, I. S., Torres, M., Ponte, C. R., Ruiz, Y., Salas, A., Nguyen, E., Eng, C., Borjas, L., … LACE Consortium. (2012). Development of a panel of genome-wide ancestry informative markers to study admixture throughout the Americas. PLoS Genetics, 8(3), e1002554.

Haller, B. C., & Messer, P. W. (2023). SLiM 4: Multispecies Eco-Evolutionary Modeling. The American Naturalist, 201(5), E127–E139.

Hess, J. E., Campbell, N. R., Docker, M. F., Baker, C., Jackson, A., Lampman, R., McIlraith, B., Moser, M. L., Statler, D. P., Young, W. P., Wildbill, A. J., & Narum, S. R. (2015). Use of genotyping by sequencing data to develop a high-throughput and multifunctional SNP panel for conservation applications in Pacific lamprey. Molecular Ecology Resources, 15(1), 187–202.

Hoban, S., Gaggiotti, O., Bertorelle, G., & ConGRESS Consortium. (2013). Sample Planning Optimization Tool for conservation and population Genetics (SPOTG): a software for choosing the appropriate number of markers and samples. Methods in Ecology and Evolution / British Ecological Society, 4(3), 299–303.

Kechin, A., Borobova, V., Boyarskikh, U., Khrapov, E., Subbotin, S., & Filipenko, M. (2020). NGS-PrimerPlex: High-throughput primer design for multiplex polymerase chain reactions. PLoS Computational Biology, 16(12), e1008468.

Kidd, K. K., Pakstis, A. J., Speed, W. C., Grigorenko, E. L., Kajuna, S. L. B., Karoma, N. J., Kungulilo, S., Kim, J.-J., Lu, R.-B., Odunsi, A., Okonofua, F., Parnas, J., Schulz, L. O., Zhukova, O. V., & Kidd, J. R. (2006). Developing a SNP panel for forensic identification of individuals. Forensic Science International, 164(1), 20–32.

Kleinman-Ruiz, D., Martínez-Cruz, B., Soriano, L., Lucena-Perez, M., Cruz, F., Villanueva, B., Fernández, J., & Godoy, J. A. (2017). Novel efficient genome-wide SNP panels for the conservation of the highly endangered Iberian lynx. BMC Genomics, 18(1), 556.

Manel, S., Gaggiotti, O. E., & Waples, R. S. (2005). Assignment methods: matching biological questions with appropriate techniques. Trends in Ecology & Evolution, 20(3), 136–142.

McVean, G. (2009). A genealogical interpretation of principal components analysis. PLoS Genetics, 5(10), e1000686.

Mooney, J. A., Marsden, C. D., Yohannes, A., Wayne, R. K., & Lohmueller, K. E. (2023). Longterm Small Population Size, Deleterious Variation, and Altitude Adaptation in the Ethiopian Wolf, a Severely Endangered Canid. Molecular Biology and Evolution, 40(1). 10.1093/molbev/msac277

Mooney, J. A., Yohannes, A., & Lohmueller, K. E. (2021). The impact of identity by descent on fitness and disease in dogs. Proceedings of the National Academy of Sciences, 118(16), e2019116118.

Morin, P. A., Martien, K. K., & Taylor, B. L. (2009). Assessing statistical power of SNPs for population structure and conservation studies. Molecular Ecology Resources, 9(1), 66–73.

Morrell, P. L., & Clegg, M. T. (2007). Genetic evidence for a second domestication of barley (Hordeum vulgare) east of the Fertile Crescent. Proceedings of the National Academy of Sciences of the United States of America, 104(9), 3289–3294.

Natesh, M., Taylor, R. W., Truelove, N. K., Hadly, E. A., Palumbi, S. R., Petrov, D. A., & Ramakrishnan, U. (2019). Empowering conservation practice with efficient and economical genotyping from poor quality samples. Methods in Ecology and Evolution / British Ecological Society, 10(6), 853–859.

Nicholson, G., Smith, A. V., Jónsson, F., Gústafsson, Ó., Stefánsson, K., & Donnelly, P. (2002). Assessing Population Differentiation and Isolation from Single-Nucleotide Polymorphism Data. Journal of the Royal Statistical Society. Series B, Statistical Methodology, 64(4), 695–715.

Oliveira, R., Randi, E., Mattucci, F., Kurushima, J. D., Lyons, L. A., & Alves, P. C. (2015). Toward a genome-wide approach for detecting hybrids: informative SNPs to detect introgression between domestic cats and European wildcats (Felis silvestris). Heredity, 115(3), 195–205.

Pakstis, A. J., Speed, W. C., Fang, R., Hyland, F. C. L., Furtado, M. R., Kidd, J. R., & Kidd, K. K. (2010). SNPs for a universal individual identification panel. Human Genetics, 127(3), 315–324.

Pakstis, A. J., Speed, W. C., Kidd, J. R., & Kidd, K. K. (2007). Candidate SNPs for a universal individual identification panel. Human Genetics, 121(3-4), 305–317.

Parker, H. G., Dreger, D. L., Rimbault, M., Davis, B. W., Mullen, A. B., Carpintero-Ramirez, G., & Ostrander, E. A. (2017). Genomic Analyses Reveal the Influence of Geographic Origin, Migration, and Hybridization on Modern Dog Breed Development. Cell Reports, 19(4), 697–708.

Parker, L. D., Campana, M. G., Quinta, J. D., Cypher, B., Rivera, I., Fleischer, R. C., Ralls, K., Wilbert, T. R., Boarman, R., Boarman, W. I., & Maldonado, J. E. (2022). An efficient method for simultaneous species, individual, and sex identification via in-solution single nucleotide polymorphism capture from low-quality scat samples. Molecular Ecology Resources, 22(4), 1345–1361.

Paschou, P., Ziv, E., Burchard, E. G., Choudhry, S., Rodriguez-Cintron, W., Mahoney, M. W., & Drineas, P. (2007). PCA-correlated SNPs for structure identification in worldwide human populations. PLoS Genetics, 3(9), 1672–1686.

Patterson, N., Price, A. L., & Reich, D. (2006). Population structure and eigenanalysis. PLoS Genetics, 2(12), e190.

Pearman, W. S., Urban, L., & Alexander, A. (2022). Commonly used Hardy-Weinberg equilibrium filtering schemes impact population structure inferences using RADseq data. Molecular Ecology Resources, 22(7), 2599–2613.

Pfaff, C. L., Barnholtz-Sloan, J., Wagner, J. K., & Long, J. C. (2004). Information on ancestry from genetic markers. Genetic Epidemiology, 26(4), 305–315.

Pfaffelhuber, P., Grundner-Culemann, F., Lipphardt, V., & Baumdicker, F. (2020). How to choose sets of ancestry informative markers: A supervised feature selection approach. Forensic Science International. Genetics, 46, 102259.

Puckett, E. E. (2017). Variability in total project and per sample genotyping costs under varying study designs including with microsatellites or SNPs to answer conservation genetic questions. Conservation Genetics Resources, 9(2), 289–304.

Ramachandran, S., Deshpande, O., Roseman, C. C., Rosenberg, N. A., Feldman, M. W., & Cavalli-Sforza, L. L. (2005). Support from the relationship of genetic and geographic distance in human populations for a serial founder effect originating in Africa. Proceedings of the National Academy of Sciences of the United States of America, 102(44), 15942–15947.

Rosenberg, N. A. (2005). Algorithms for selecting informative marker panels for population assignment. Journal of Computational Biology: A Journal of Computational Molecular Cell Biology, 12(9), 1183–1201.

Rosenberg, N. A., Li, L. M., Ward, R., & Pritchard, J. K. (2003). Informativeness of genetic markers for inference of ancestry. American Journal of Human Genetics, 73(6), 1402–1422.

Rosenberg, N. A., Mahajan, S., Ramachandran, S., Zhao, C., Pritchard, J. K., & Feldman, M. W. (2005). Clines, clusters, and the effect of study design on the inference of human population structure. PLoS Genetics, 1(6), e70.

Sottile, G., Sardina, M. T., Mastrangelo, S., Di Gerlando, R., Tolone, M., Chiodi, M., & Portolano, B. (2018). Penalized classification for optimal statistical selection of markers from high-throughput genotyping: application in sheep breeds. Animal: An International Journal of Animal Bioscience, 12(6), 1118–1125.

Storer, C. G., Pascal, C. E., Roberts, S. B., Templin, W. D., Seeb, L. W., & Seeb, J. E. (2012). Rank and order: evaluating the performance of SNPs for individual assignment in a nonmodel organism. PloS One, 7(11), e49018.

Wehrenberg, G., Tokarska, M., Cocchiararo, B., & Nowak, C. (2024). A reduced SNP panel optimised for non-invasive genetic assessment of a genetically impoverished conservation icon, the European bison. Scientific Reports, 14(1), 1875.

Wilkinson, S., Wiener, P., Archibald, A. L., Law, A., Schnabel, R. D., McKay, S. D., Taylor, J. F., & Ogden, R. (2011). Evaluation of approaches for identifying population informative markers from high density SNP chips. BMC Genetics, 12, 45.

Willing, E.-M., Dreyer, C., & van Oosterhout, C. (2012). Estimates of Genetic Differentiation Measured by FST Do Not Necessarily Require Large Sample Sizes When Using Many SNP Markers. PloS One, 7(8), e42649.

Wright, S. (1951). The genetical structure of populations. Annals of Eugenics, 15(4), 323–354.

