## SupplementaryFigures for "Recommendations for Population and Individual Diagnostic SNP Selection in Non-Model Species"

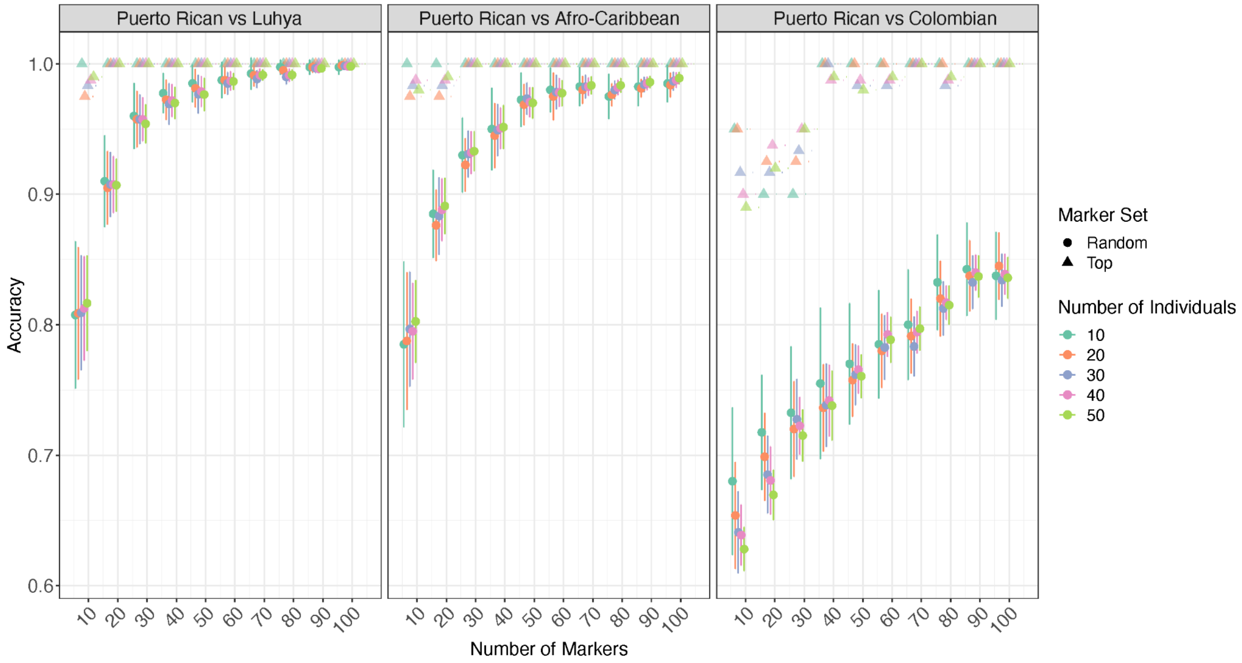


**Supplementary Figure 1:** Classification accuracy using two approaches, the top markers (triangles) and random markers (circles). For the random markers, each dot signifies the mean over 20 replicates. The x-axis indicates the number of markers used for classification and the dot color indicates the number of individuals. The accuracy of classification is shown on the y-axis. Here, three classification was conducted using Puerto Rican and Luhya, Puerto Rican and Afro-Caribbean, and Puerto Rican and Colombian.

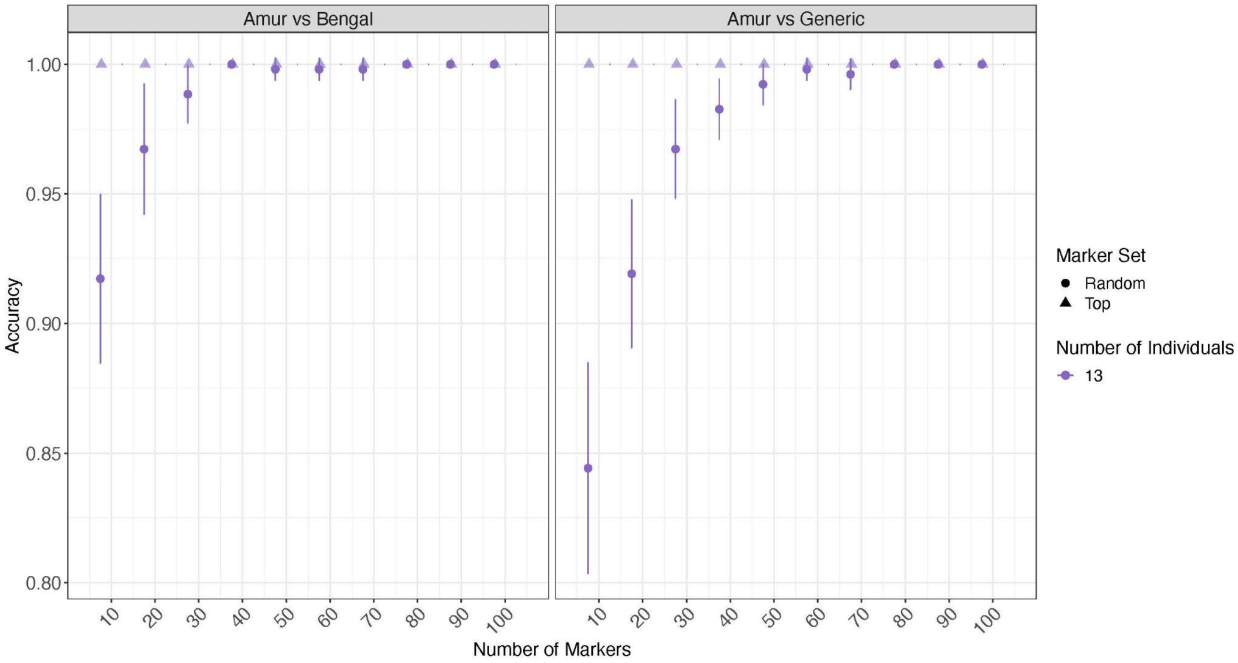


**Supplementary Figure 2:** Classification accuracy using two approaches, the top markers (triangles) and random markers (circles). For the random markers, each dot signifies the mean over 20 replicates. The x-axis indicates the number of markers used for classification and the dot color indicates the number of individuals. The accuracy of classification is shown on the y-axis. Here, two classification was conducted using Amur and Bengal tigers and Amur and Generic tigers.

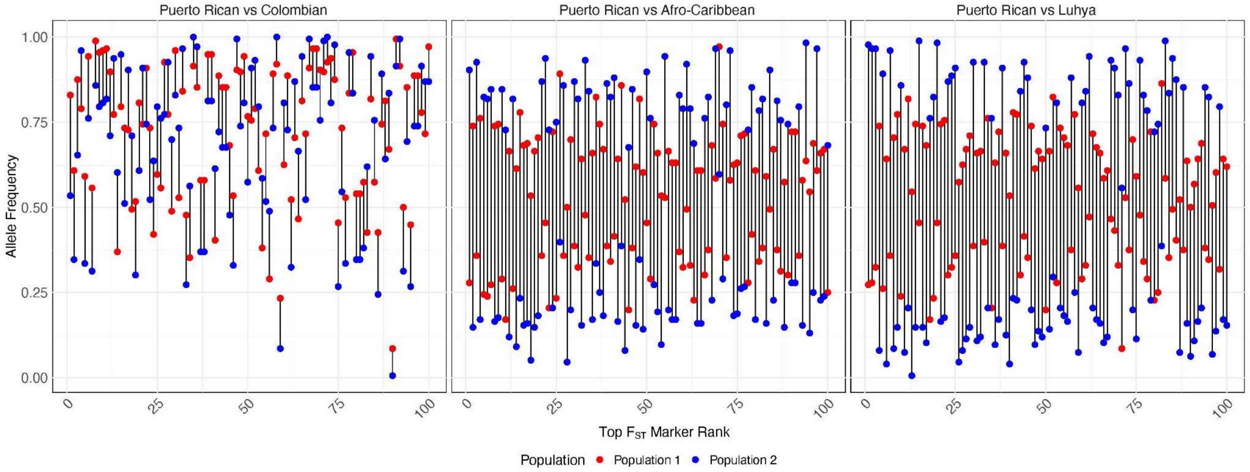


**Supplementary Figure 3:** Allele frequency of top 100 $F_{ST}$ markers between populations and the frequency difference for each marker. The x-axis indicates the rank of $F_{ST}$ values of the markers, and the y-axis indicates the markers' allele frequency in both Puerto Rican (represented in blue) and Colombian (represented in red) populations; Puerto Rican and Afro-Caribbean (represented in red) populations; and Puerto Rican and Luhya (represented in red) populations.

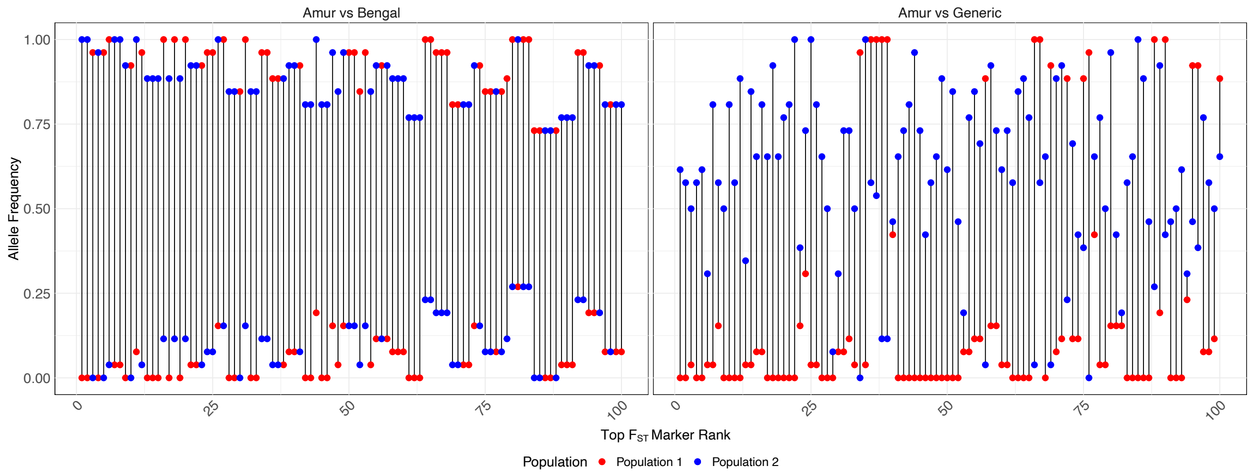


**Supplementary Figure 4:** Allele frequency of top 100 $F_{ST}$ markers between populations and the frequency difference for each marker. The x-axis indicates the rank of $F_{ST}$ values of the markers, and the y-axis indicates the markers' allele frequency in both Amur (represented in blue) and Bengal (represented in red) and on the right Amur and Generic (represented in red).

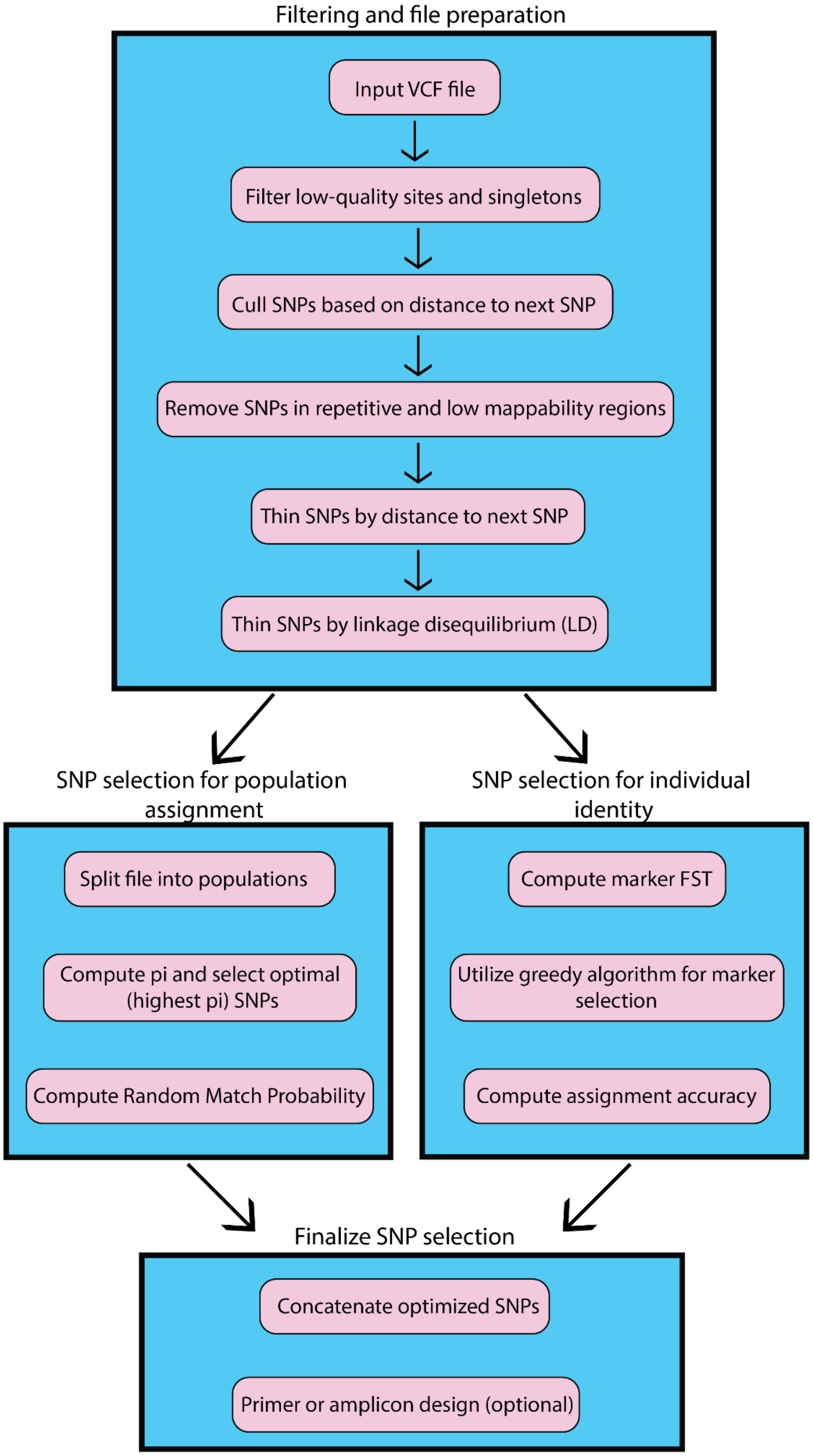


**Supplementary Figure 5**: Simplified flow chart of mPCRselect pipeline.
